# Geometric Kinematics of Human Eyes

**DOI:** 10.64898/2026.04.10.716809

**Authors:** Jacek Turski

**Affiliations:** Department of Mathematics and Statistics University of Houston-Downtown Houston, TX 77002

**Keywords:** Asymmetric eye model, Eye’s torsion-free (geodesic) rotation, Ocular torsion, Rodrigues’ vector, Rotation matrix, Euler’s angles, Angular velocity

## Abstract

In previous studies by the author on binocular vision with the asymmetric eye (AE), which models a healthy human eye with misaligned optical components, the results were primarily presented in the Rodrigues’ vector (RV) framework and supported by simulations and 3D visualizations in *GeoGebra*’s dynamic geometry environment. In this paper, the novel geometric kinematics of the human eye, that is, the eye with misaligned optics, and simplified assumptions about the eye rotations (the eye’s translational movements are disregarded), are developed within the framework of rigidbody rotations. The originality of the analysis lies in a precise geometric decomposition of a full rotation of the eye’s posture into a torsion-free rotation (the geodesic part) and a torsional rotation (the non-geodesic extension of the geodesic part). This decomposition is extended to the corresponding decomposition of the angular velocity. A novel derivation of the eye’s angular velocity from the RV formulation of the eye kinematics is proposed.

## 1 Introduction

The asymmetric eye (AE), comprising misaligned optical components in the healthy eye [1–5], was first proposed in [6]. It was developed to correct and anatomically motivate classical studies that modeled empirical horopter curves as conic sections in binocular vision using analytic geometry, first done by Ogle in 1932 [7] for symmetric fixations and later extended by Amigo in 1965 [8] to any fixations.

For a comprehensive discussion of the AE model, refer to [9, 10]. The binocular system with AEs, including horizontal misalignment, was used in [9, 11] to study the geometric properties of a family of iso-disparity curves, where zero disparity corresponds to the horopter. It is well known that the shape of the empirical horopter is the result of an asymmetry in horizontal retinal correspondence. However, the origin of this asymmetry has been uncertain [12, 13]. It was demonstrated in [11] that the distribution of iso-disparity curves (the zero-disparity curve is the horopter) was invariant with respect to lens tilt for a relatively stable displacement of the fovea in the human population. This invariance shows that the distribution of iso-disparity curves is *universal* for the eye’s optical misalignment. It established, for the first time, that the eye’s optical asymmetry implies the shape of the horopter resembling the empirical horopter rather than the commonly used Vieth-Müller circle (VMC).

Further, the global aspects of visual space’s phenomenal spatial relations—the variable curvature and finite horizon—were studied in [11] for the binocular system with AEs in the framework of Riemannian geometry. The global aspects of stereopsis are crucial to perception, as exemplified by the coarse disparity contribution to our impression of being immersed in the ambient environment despite receiving 2D retinal projections of light reflected by spatial objects [14, 15].

In [10], the vertical misalignment of the eye’s optical components was added to complement the horizontal misalignment. As discussed in [10], the lens’s vertical misalignment accounted for the backward tilt of the vertical horopter seen by observers. It replaced Helmholtz’s classic theory, which held that the inclination of the subjective vertical retinal meridian to the retinal horizon explains the backward tilt of the perceived vertical horopter. This replacement reflects that retinal meridians are not well defined when the eye’s optical components are misaligned. Furthermore, the large declination angles between the subjective vertical meridians of the right and left eyes, as reported by Helmholtz and others, are ‘idiosyncratic’ (see the discussion in [16]) and therefore cannot be accepted as representative of the average normal response. In contrast, the criterion of retinal vertical correspondence based on the vertical lens tilt accounts for the observed vertical horopter inclination relative to the true vertical in more recent experiments [16, 17].

By considering the common human eye’s misaligned optics, ab initio binocular Listing’s law, and the half-angle rule, we formulate them in [18] using Rodrigues’ vector, simulate and visualize them in *GeoGebra*, and support them with modern ophthalmology studies. Listing’s law constrains the eye’s torsional degrees of freedom by assuming that for the eye’s primary position, the eye’s rotation vector is contained in the Listing plane. The half-angle rule extends Listing’s law to changes in the fixation axis between the eye’s tertiary positions. As already discussed in [10], in the binocular formulation of Listing’s law, the primary eye position is replaced by the binocular eyes’ resting posture (ERP) corresponding to the eye muscles’ natural tonus resting position, which serves as a zero-reference level for convergence effort. It resolved two outstanding issues: the lack of a generally accepted explanation for Listing’s law [19] and the elusive neurophysiological significance of the primary position and the Listing plane [20].

The main original results of [18] include constructing the configuration space of sequences of 3D changes in the bifixating eyes. The configuration space construction, combined with simulations, demonstrates the binocular half-angle rule. Because of misaligned-eye optics, the optical axis, the visual axis, the lens’s optical axis, and the fixation axis differ, as shown in Fig. 1. This makes the geometric definition of ocular torsion intricate. The results in [18] were obtained by decomposing each bifixating eye’s positional change into a torsion-free rotation (geodesic part) that changes the visual axis direction and a torsional rotation (non-geodesic extension) that best approximates rotation about the lens’ optical axis.

**Figure 1.**
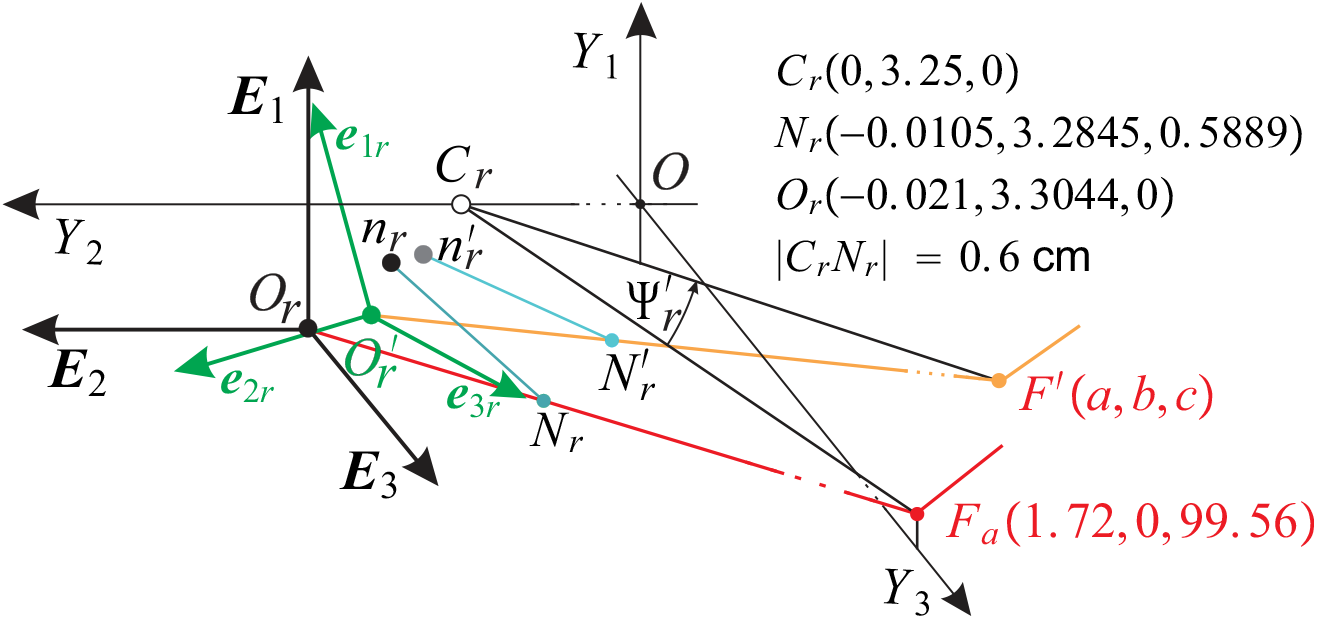
The right AE. In the ERP, each AE’s image plane is coplanar with the head frontal *Y*_1_*Y*_2_-plane. For the right AE, this image plane is spanned by the vectors ***E***_1_ and ***E***_2_ of the black frame {***E***_1_, ***E***_2_, ***E***_3_} attached at optical center *O*_*r*_, the projection along the (red) visual axis passing through the ERP fixation *F*_*a*_ and the nodal point *N*_*r*_. The vector ***E***_3_ is parallel to the lens (blue) optical axis through *n*_*r*_ and *N*_*r*_. The right AE rigidly rotates with the image plane when the eyes’ rotation at *C*_*r*_ changes the fixation axis through *F*_*a*_ to a fixation axis through *F*^*′*^. The green frame {***e***_1*r*_, ***e***_2*r*_, ***e***_3*r*_} at 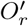, results from the AE rotation at *C*_*r*_. The rotated plane is spanned by vectors ***e***_1*r*_ and ***e***_2*r*_. The green frame is translated by 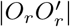 with the corresponding translated points 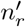 and 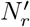, which are not included in the geometric analysis, as explained in the text. The coordinates of *C*_*r*_, *N*_*r*_, and *O*_*r*_ express the anthropomorphic eye’s dimensions.

In this work, I extend prior studies on 3D positional changes in the binocular system with AEs to 3D rotation kinematics. The geometric parametrization of the AE positional change in [18] is modified here to be compatible with ‘*z*-*x*-*z*’ original Euler-angle parametrization used frequently in the analysis of rigid body rotations, see [21], for example. We carefully study the geometry of the eyes’ kinematics decomposition into torsion-free and torsional parts, similar to the parametrization first proposed by Piña in [22] in the mathematical treatment of rigid-body dynamics. A novel to Piña’s derivation of the eye’s angular velocity from the RV formulation of the eye kinematics is proposed. The corresponding decomposition of the angular velocity is discussed.

## 2 Review of the Binocular System with AEs

### 2.1 Asymmetry angles in AE and the ERP

The typical asymmetry angles in the human population for the fovea’s horizontal and vertical displacements from the posterior pole are *α* = 5.2°and *γ* = −2°and the lens’ horizontal and vertical tilts relative to the optical axis are *β* = 3.3°and *ε* = −1°. For the definition of the misalignment angles in the AE model, refer to Fig. 2 in [10], where the angles are shown for the right eye.

**Figure 2.**
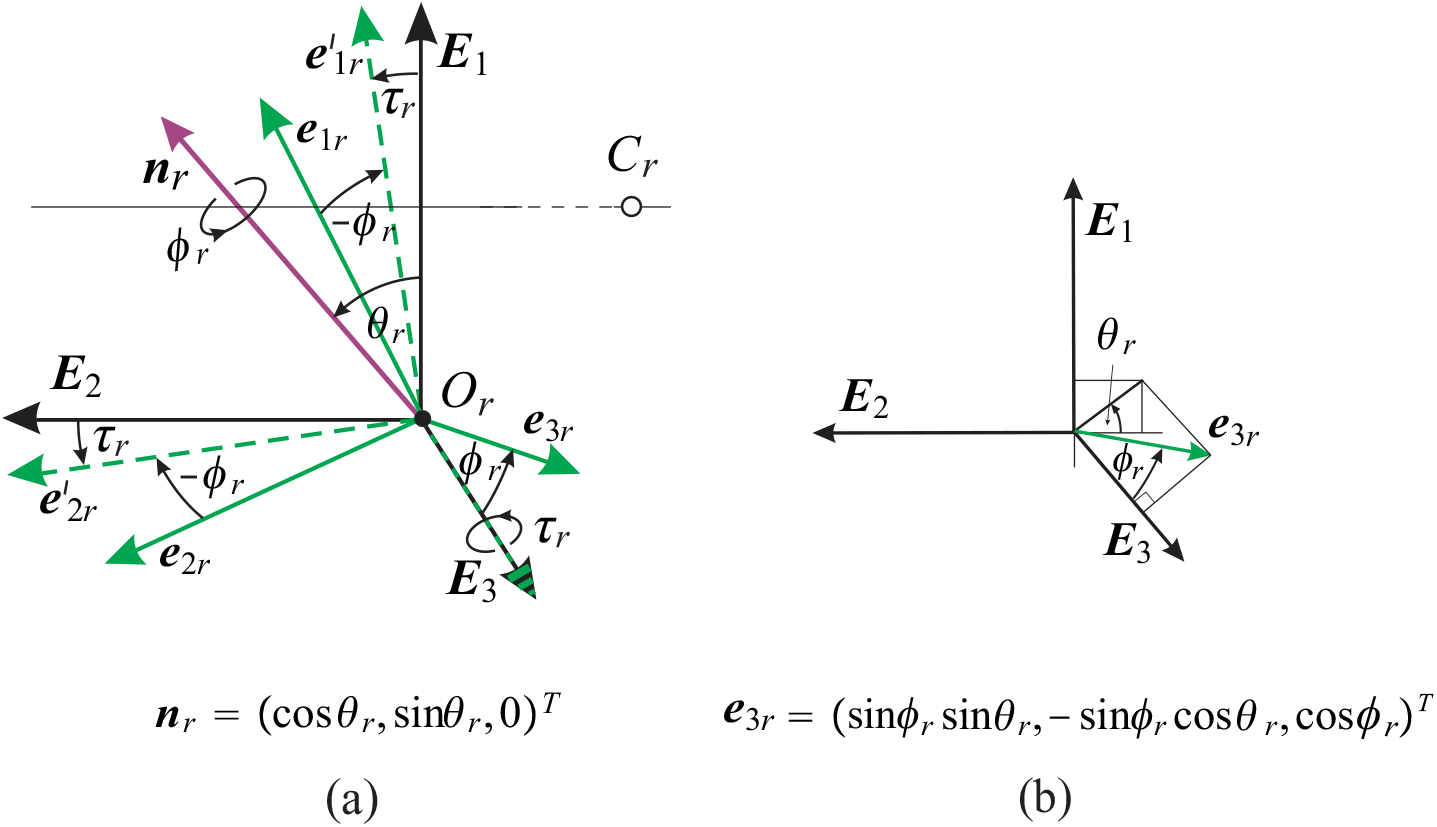
The right AE view is shown in the ERP near the optical center *O*_*r*_. (a) The AE image-plane orientation is specified by the frame vectors ***E***_1_ and ***E***_2_. After AE is rotated at *C*_*r*_, the frame {***E***_1_, ***E***_2_, ***E***_3_} changes to {***e***_1*r*_, ***e***_2*r*_, ***e***_3*r*_}. RV ***r***_*r*_ = tan(*ϕ*_*r*_*/*2)***n***_*r*_, where ***n***_*r*_ = (cos *θ*_*r*_, sin *θ*_*r*_, 0)^*T*^ rotates ***E***_3_ onto ***e***_3*r*_ in the plane spanned by these vectors. Under rotation by −***r***_*r*_, the frame {***e***_1*r*_, ***e***_2*r*_, ***e***_3*r*_} changes to 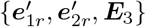. Ocular torsion is *τ*_*r*_. (b) The unit vector ***e***_3*r*_ perpendicular to the right eye rotated image plane (spanned by ***e***_1*r*_ and ***e***_2*r*_ shown in (a)) is expressed in terms of the angles *ϕ*_*r*_ and *θ*_*r*_. The details are given in the text.

The binocular system with the AEs has a unique bifixation for the given asymmetry angles, such that the resulting eye’s ERP is specified by the lenses’ coplanar equatorial planes being parallel to the coplanar image planes aligned with the head’s frontal plane. For the values of asymmetry angles listed above, the fixation of the ERP is *F*_*a*_(1.72, 0, 99.56), with coordinates in centimeters matching the average abathic distance of about 1 m of the empirical horopter distinguished by its straight, frontal line contained in the coplanar image planes; see Fig. 5 in [10].

In Fig. 1, we changed the coordinate orientation from previous studies to align with the description in [21, 23]. In the ERP, the right AE’s coordinates at *O*_*r*_—the optical center on the image plane obtained by projecting the fovea through the nodal point along the red visual axis connecting *F*_*a*_ and *O*_*r*_ in Fig. 1—are given by the frame {***E***_1_, ***E***_2_, ***E***_3_}. The left AE frame consists of the same vectors placed at *O*_*l*_, which is mirror-symmetric to *O*_*r*_ in the mid-sagittal plane (the *Y*_1_*Y*_3_-plane shown in Fig.1). We see that, ***E***_1_ = (1, 0, 0)^*T*^, ***E***_2_ = (0, 1, 0)^*T*^ and ***E***_3_ = (0, 0, 1)^*T*^ provides coordinates (*X*_1*r*_, *X*_2*r*_, *X*_3*r*_) at *O*_*r*_ for the right AE and with coordinates (*X*_1*l*_, *X*_2*l*_, *X*_3*l*_) at *O*_*l*_ for the left AE. Both sets of coordinates are parallel to the head coordinates (*Y*_1_, *Y*_2_, *Y*_3_) at *O*. The vector ***E***_3_ is parallel to the lenses’ optical axes through *n*_*r*_ and *N*_*r*_, where *N*_*r*_ is the right nodal point.

For the values of asymmetry angles listed above, the fixation of the ERP is *F*_*a*_(1.72, 0, 99.56), with coordinates in centimeters matching the average abathic distance of about 1 m of the empirical horopter distinguished by its straight, frontal line contained in the co-planar image planes; see Fig. 5 in [10]. Fig. 2 in cite turski2025 discusses the difference between the schematic eye (symmetric, single refractive eye model) and the AE and its impact on the horopteric curves. The main difference is that the horopters in the binocular system with AEs resemble the empirical horopters, whereas the horopters in the binocular system with the schematic eyes are the Vieth-Müller circles, approximating the empirical horopters near the fixation point and significantly deviating farther distance along the horopter from the fixation point.

### 2.2 Orientation change in AEs

Fig. ref fig:1 schematically shows the geometry of the orientation change of the AEs in the right eye when the bifixation changes from *F*_*a*_ to *F*^*′*^. The orientation change of the AE is produced by rotating the fixation axes at *C*_*r*_, i.e., the rotation of the line through *C*_*r*_ and *F*_*a*_ into the line through *C*_*r*_ and *F*^*′*^. These rotations follow the Euler theorem for each eye; each rotation is about a line passing through the rotation center, cf. [21].

Because the eye’s gaze change is produced by the fixation axis rotation at *C*_*r*_, the EA change in orientation is given by the resulting change of the frame {***E***_1_, ***E***_2_, ***E***_3_} attached at the optical center *O*_*r*_ of the image plane into the green frame {***e***_1*r*_, ***e***_2*r*_, ***e***_3*r*_} attached at 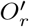. The corresponding discussion holds for the left AE, that is, the frame {***E***_1_, ***E***_2_, ***E***_3_} attached at the optical center *O*_*l*_ of the image plane into the frame e_*l*_ = {***e***_1*l*_, ***e***_2*l*_, ***e***_3*l*_} attached at 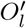.

From the anthropometric data presented in Fig. 1, the translation lengths from *O*_*r*_ to 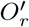 and from *O*_*r*_ to 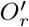 are less than 0.03 cm for a typical eye’s rotation angles, which is much smaller than the distances to the fixation points. Although this translation is not included in the geometric analysis presented here, it has always been included in *GeoGebra* simulations of Listing’s law in [10, 18]. Furthermore, the eyeball is suspended in the fatty layer of the bony orbit and, hence, under the action of six extraocular muscles, undergoes a slight translational motion estimated at about 0.06 cm [24] in addition to the main rotational motion. This translation is also not considered in this study.

## 3 AEs bifixation changes with Rodrigues vectors

Rodrigues’ (rotation) vectors (RVs) efficiently represent 3D rotations, encoding the rotation axis and angle with three parameters—the minimum required. As shown in [18], RVs are particularly well-suited for discussing Listing’s law and the half-angle rule, which posits that the change in eye position is given by the rotation vector in the displacement plane determined by the reference fixation. When the eyes’ positions change from the ERP fixation, the RV lies in the image plane, which coincides with the stationary upright head’s frontal plane and is parallel to the lens equatorial plane; the binocular Listing’s law is formulated in [18]. When the eyes bifixate at the tertiary position, RV lies in the ERP image plane, tilted by half the eye’s eccentricity angle; the binocular half-angle rule extends the binocular Listing’s law.

### 3.1 Definition of Rodrigues’ vector

To introduce the RV framework, I start with the conclusion from Euler’s rotation theorem: Any rotation matrix *R* can be parametrized as *R*(*φ*, ***n***) for a rotation angle *φ* around the axis with the unit column vector ***n***. This parametrization is unique if you fix the orientation to 0 *< φ <* 180°. Usually, a counterclockwise (or right-hand) orientation is used to define the positive values of angles.

The pair *ρ* = cos(*φ/*2), ***e*** = sin(*φ/*2)***n*** is known as *Euler-Rodrigues parameters* and

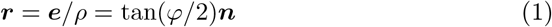

as *RV*, usually referred to as *rotation vector* [23]. Rodrigues proved that under composition *R*(*φ*, ***n***) = *R*(*φ*^*′′*^, ***n***^*′′*^)*R*(*φ*^*′*^, ***n***^*′*^), the corresponding Euler-Rodrigues parameters transform as follows [25],

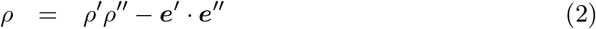

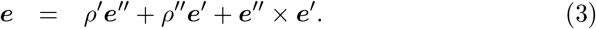

Then, setting ***r*** = ***e****/ρ*, we see that the composition for RVs is given by

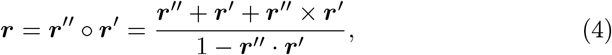

Further, ***r***^*−*1^ = −***r*** and tan(−*φ/*2)***n*** = tan(*φ/*2)(−***n***) define the same rotation vector. As is shown in (4), the parallelogram law does not hold for RVs because rotations are not commutative.

Note that (cos(*φ/*2), sin(*φ/*2)**n**) is a unit quaternion that describes the rotation by *ϕ* around **n**. Westheimer introduced quaternions in 1957 [26] to describe eye kinematics. Quaternions and Rodrigues’ (rotation) vectors have often been used to analyze the eye’s rotations with the connection to Listing’s law, cf. [27–31].

### 3.2 RV and rotation matrix

Following [23], the rotation matrix can be expressed in terms of RV *r* as

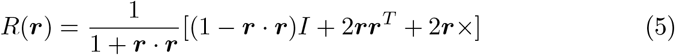

where for the column vector **r** = (*r*_1_, *r*_2_, *r*_3_)^*T*^ (“T” denotes the transpose) and

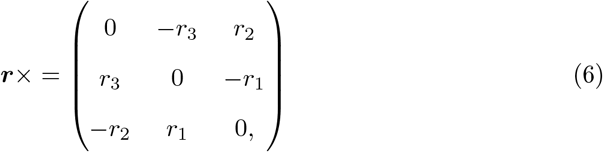

such that the matrix ***r***× at (6) acts on a vector ***v*** giving the familiar cross product: **(*r***×)***v*** = ***r*** × ***v***. Further,

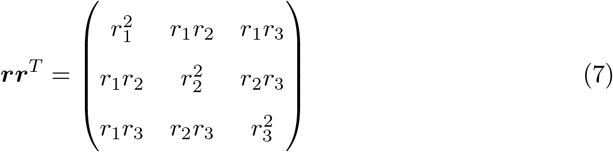

so that ***rr***^*T*^ ***v*** = (***r*** · ***v***)***r***.

For completeness of discussion, the rotation matrix (5) acts on a vector ***v*** as follows

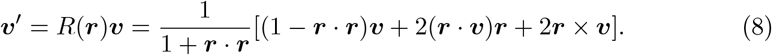

## 4 Simulation of the Half-angle rule with RVs

## 5 Torsion-free and torsional components in AE change of position

Torsion-free and torsional rotation are strictly defined for the eyes. Torsion-free rotation changes the direction of the eye’s visual axis, whereas torsional rotation changes the eye’s orientation about the lens optical axis. In vision science, the reduced, symmetric eye (SE) model is typically used, in which the visual axis coincides with the lens optical axis. Because I use the AE here to model the inherent misalignment of the human eye’s optical components, the visual axis differs from the lens optical axis, so these rotations are approximate yet highly accurate. The reader is referred to [18], where the differences between SE and AE are discussed.

### 5.1 Geometry of the AE frame rotation

As shown in Fig. 1, changes in the right AE’s orientation are expressed in terms of frame vectors attached at the optical center *O*_*r*_, which is given by the projection of the fovea through the nodal point into the image plane along the visual axis (the red line in Fig. 1) coming from the part of the object being looked at (fixation point). It is the path of light for clearest vision. The visual axis is different from the lens’s optical axis (the blue line in Fig. 1). It also differs from the fixation axis, which connects the eye’s rotation center to the fixation point. This ubiquitous misalignment of the human eye’s optical components is included in the first geometric discussion of eye rotations.

This section discusses the geometry of eye rotations in the binocular system with AEs, emphasizing the decomposition into torsion-free rotations and torsional rotations. Ocular torsion is important for clinical diagnosis [32] and is often measured using video-oculography, a non-invasive, high-resolution technique. A robust method for measuring ocular torsion using this technique was recently proposed in [33], which tracks the angular shift of the iris pattern about the pupil center and offers advantages over previous methods.

[18] considered the parametrization of AE rotations, supported by simulations and 3D visualization in *GeoGebra*’s dynamic geometry environment. Here, I investigate its geometric formulation for one eye by decomposing it into a torsionfree rotation, which describes changes in the direction of the visual axis, and a torsional rotation, which best approximates ocular torsion about the lens optical axis. This work will be crucial for future studies of the AEs’ dynamic description, which involves the oculomotor plant but is outside the scope of this study. Crucially, this decomposition enables straightforward derivation of the rotation matrix and discussion of angular rotation.

Referring to Fig. 2, I first define for the right AE the rotation matrix *R*_*r*_(*ϕ*_*r*_, ***n***_*r*_) that rotates ***E***_3_ to ***e***_3*r*_ in the plane spanned by these two vectors, *R*(*ϕ*_*r*_, ***n***_*r*_)***E***_3_ = ***e***_3*r*_. The inverse rotation, when applied to the whole frame, is the following: *R*(−*ϕ*_*r*_, ***n***_*r*_)***e***_3*r*_ = ***E***_3_ and 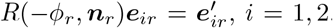. Thus, 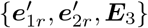 is an orthogonal frame, which justifies the definition of the torsional angle *τ*_*r*_ by 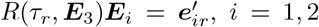, *i* = 1, 2, shown in Fig. 2. It implies that 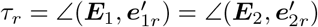 expresses the ocular torsion. Because the torsional rotations are about ***E***_3_, which is parallel to the lens optical axis at a distance of 0.02 cm, *τ*_*r*_ best approximates rotation about the lens optical axis compared with other axes (visual and fixation axes). In vision science, ocular torsion is defined as rotation about the visual axis.

Then, we can check that

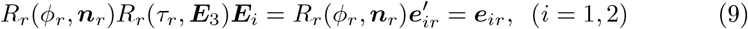

such that we have

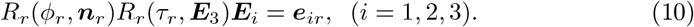

### 5.2 RVs of torsional and geodesic rotations

The torsional angle *τ*_*r*_ for the rotation from the ERP defines the corresponding RV by

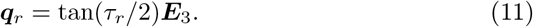

For the left AE’s, we have a similar expression ***q***_*l*_ = tan(*τ*_*l*_*/*2)***E***_3_.

Next, the rotation represented by RV

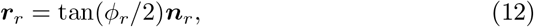

which rotates ***E***_3_ to ***e***_3*r*_ in the plane perpendicular to these vectors gives the shortest path on the group *SO*(3) that takes ***E***_3_ to ***e***_3*r*_ along the great circle of the sphere centered at *O*_*r*_, i.e., the geodesic [34]. The rotation represented by ***r***_*r*_ °***q***_*r*_ is no longer a geodesic, and ***q***_*r*_ describes how far the composite rotation differs from a geodesic. Again, the left RV is ***r***_*l*_ = tan(*ϕ*_*l*_*/*2)***n***_*l*_, ***q***_*l*_ = tan(*τ*_*l*_*/*2)***E***_3_, and the above discussion also holds for the left eye.

I get the RVs ***r***_*r*_ = tan(*ϕ*_*r*_*/*2)***n***_*r*_ by first specifying coordinates of the vector ***e***_3*r*_ by the angles shown in Fig. 2 (b),

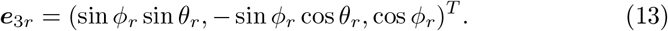

This vector is expressed in the polar coordinates of the unit sphere with azimuthal angle 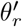 taken from ***E***_1_, i.e., 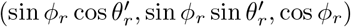 with 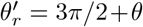. Then we see that RV ***r***_*r*_ = tan(*ϕ*_*r*_*/*2)***n***_*r*_ can be written in terms of frame vectors as

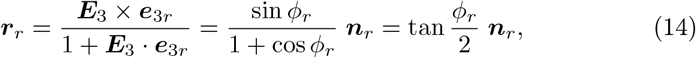

where, by performing the cross and dot products for ***e***_3*r*_ in (13) and ***E***_3_ = (0, 0, 1)^*T*^, the ***n***_*r*_ vector is as follows

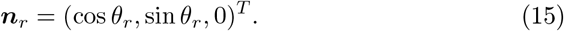

We notice from (14) and (15) the independence of the angles *θ*_*r*_ and *ϕ*_*r*_ in the definition of ***r***_*r*_; *ϕ*_*r*_ specifies the rotation angle around ***n***_*r*_ while *θ*_*r*_ specifies the direction of ***n***_*r*_. The torsion-free RV ***r***_*r*_ is complemented by the torsional RV ***q***_*r*_, given in (11) for the right AE. The corresponding expressions hold for the left AEs.

The full rotation matrix *R* = *R*(*ϕ*_*r*_, ***n***_*r*_)*R*(*τ*_*r*_, ***E***_3_), which is obtained explicitly in the next section, is related to the parametrization of rotation proposed by Piña in [22]. However, in this study, the free parameter in his parametrization is ocular torsion.

## 6 Rotation matrix of AE displacement from ERP

Following Sec. 3.2, without specifying the eye to simplify the notation, the RV ***r*** = tan(*ϕ/*2)(cos *θ*, sin *θ*, 0)^*T*^, the rotation matrix *R*(***r***) is

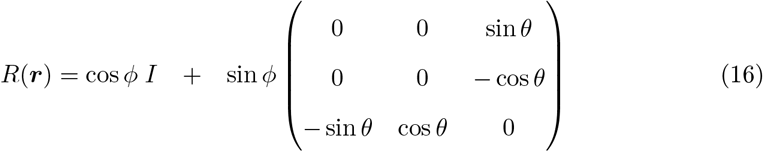

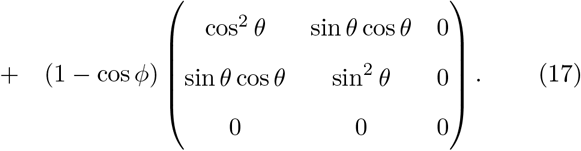

We easily get

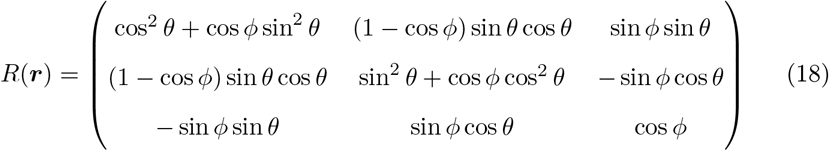

We also see that the rotation matrix for the RV ***q*** is

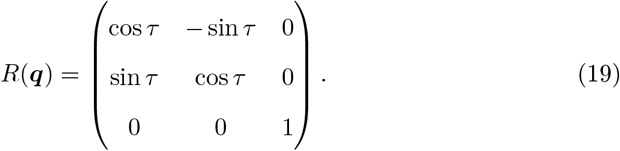

The rotation matrix for the full change in the AE position, *R*(***r***)*R*(***q***), can be calculated as follows:

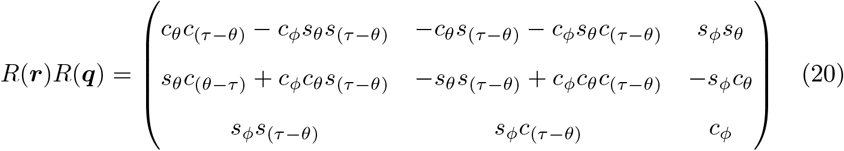

where *s*_*α*_ stands for sin *α* and *c*_*α*_ stands for cos *α*.

We note that the angle *ψ* = *τ* − *θ* appears naturally in the matrix (20). In Sec. (A.1), I show, using Euler-angle parametrization, that *τ* is the ocular torsion. Further, from the matrix in (20), we obtain the rotated frame vectors ***e***_1_, ***e***_2_, and ***e***_3_ shown in Fig. 2 that are given by the columns of this matrix as follows:

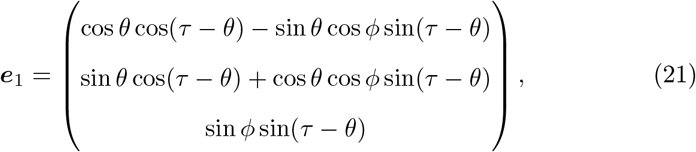

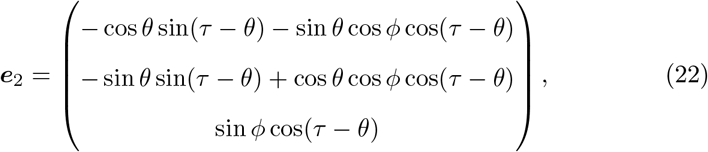

and

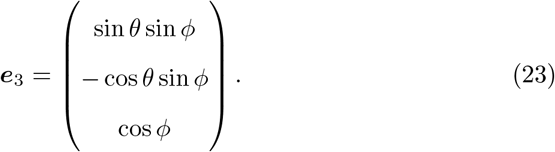

## 7 Angular velocity

### 7.1 Eye rotational movement in the ERP and in the rotating frames

In the geometric description of the eye movements, as related to the binocular Listing’s law, two frames for each AE are used: the initial ERP frames {***E***_*i*_} and the right eye rotating frames {***e***_*ir*_(*t*)} and the left eye rotating frame {***e***_*il*_(*t*)}.

The frames are placed at the optical centers on the image plane at *O*_*r*_ and *O*_*l*_ for the right and left eyes, respectively. Thus, *R*_*r*_(*t*)***E***_*i*_ = ***e***_*ir*_(*t*) for the right AE and *R*_*l*_(*t*)***E***_*i*_ = ***e***_*il*_(*t*) for the left AE.

Next, again without specifying the right or left eye, the frames are related by

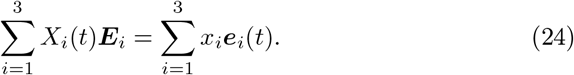

Here, *X*_*i*_(*t*) and *x*_*i*_ are coordinates of the co-rotating position vector in the initial frame and in the AE rotating frame, respectively. Then, the vector ***X***(*t*) in the basis {***E***_*i*_} and the vector ***x*** in the basis {***e***_*i*_(*t*)} can be expressed as ***X***(*t*) = *R*(*t*)***x***.

Omitting further time dependencies, the Euler-angle rotation matrix *R* = *R*_*z*_(*θ*)*R*_*x*_(*ϕ*)*R*_*z*_(*τ* − *θ*) for each eye can be expressed in terms of the rotating eye frame vectors as the matrix columns, i.e., *R* = (***e***_1_ ***e***_2_ ***e***_3_). These vectors give the precise expressions for the AEs’ rotating frames in Fig. 2.

The relation between the vector ***X*** = (*X*_1_, *X*_2_, *X*_3_)^*T*^ given in the (initial) ERP frame {***E***_*i*_} and the vector ***x*** = (*x*_1_, *x*_2_, *x*_3_)^*T*^ in the rotating eye frame {***e***_*i*_} can be written as

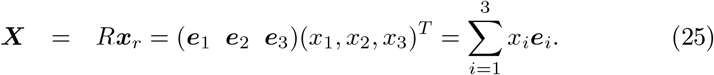

Further, because *R*^*−*1^ = *R*^*T*^,

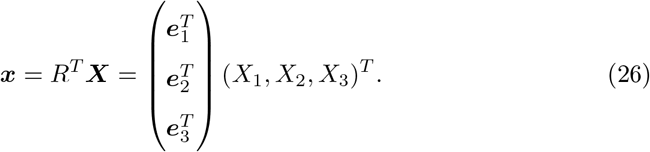

### 7.2 Angular velocity in the ERP frame and rotating frame

The angular velocity can be expressed in the eye’s moving frame or in the ERP frame and denoted by ***ω***^*ℰ*^ or by ***ω***^*I*^, respectively. The angular velocity is first derived in the ERP frame by differentiating ***X*** = *R****x*** to obtain 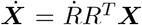.

Then, by differentiating the identity *RR*^*T*^ = *I*, we have 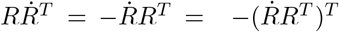 so that 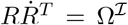 is asymmetric, which means that Ω^*I*^ = ***ω***^*I*^× as in (6). Thus, we derive 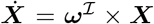. To get the angular velocity in the eye-moving frame, I note that 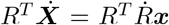. Since 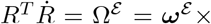, we obtain 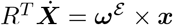.

Because the Euler parametrization conforms to the geometry of AE rotation shown in Fig. 2 for the right eye, it is possible to derive relations between the angular velocity and the rate of change of the Eulerian angles using this figure; cf., for example, Sec. 4.4 of Ch. 4 in [21] for the general discussion of rotating rigid bodies. However, the rotation matrix *R* in this study corresponds to the transpose matrix *Ã* in [21]. Since in Fig. 2, the line of nodes is given by ***n***_*r*_ = (cos *θ*, sin *θ*, 0)^*T*^ about which ***E***_3_ is rotated into ***e***_3*r*_ = (sin *ϕ* sin *θ*, − sin *ϕ* cos *θ*, cos *ϕ*)^*T*^, the angular velocity in the eye’s moving frame is

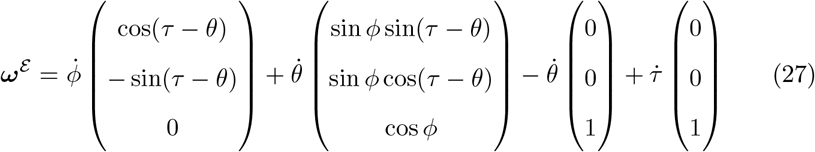

and in the ERP’s initial frame is

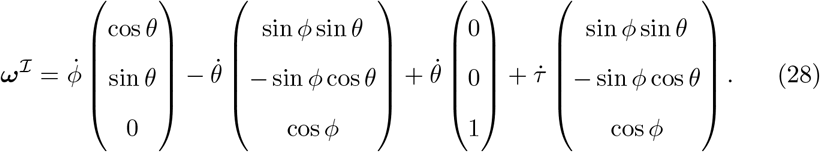

As is well known, ***ω***^*E*^ = *R*^*T*^ ***ω***^*I*^, which can also be obtained from the rotation-matrix relations given below.

In the rest of the paper, I use the following results involving the matrices *R*(***r***) and *R*(***q***) given in (18) and (19), respectively. I recall that here RVs are ***r*** = tan(*ϕ/*2)***n*** and ***q*** = tan(*τ/*2)***E***_3_.

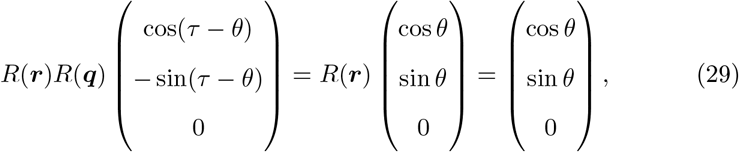

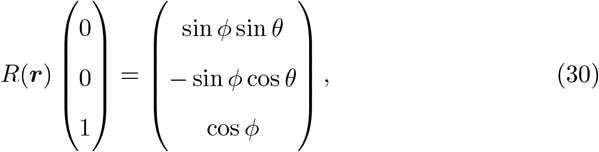

and

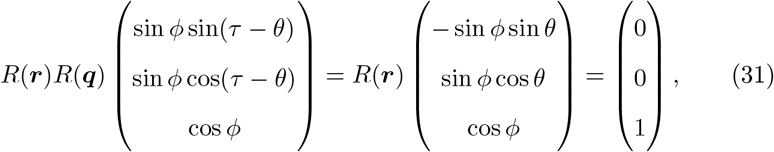

These formulas follow from aligning the frame rotations shown in Fig. 3 with the Euler-angle parametrization in (53). I note that the second equality in (29) holds because ***n*** = (cos *θ*, sin *θ*, 0)^*T*^ is the rotation axis of *R*(***r***). These results are similar to those given in [22], except that they are adapted to the AE rotation splitting into torsion-free and torsional parts.

This splitting, which was crucial for discussing the half-angle rule and the configuration space of the AEs bifixating sequences in [18], is adapted here to the angular velocity. Using (29), (30), and (31), we can write the angular velocity in torsion-free and torsional parts

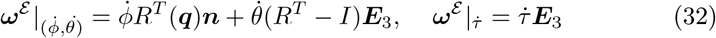

and

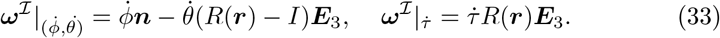

The angular velocity computed in Appendix A.1 directly from the RVs ***r*** and ***q*** (without their composition) is in the eye frame

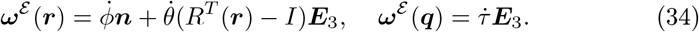

and in the ERP initial frame

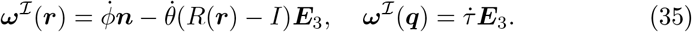

We immediately see that the relations between these different definitions of the angular velocity are

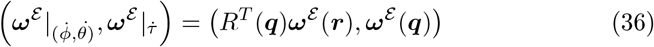

and

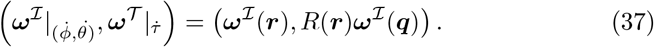

## 8 New derivation of the angular velocity

In this section, I outline the easiest way to derive the angular velocities of the RVs ***r, ρ***, and ***r*** °***ρ***. This derivation connects the composition of RVs between infinitesimal binocularly constrained tertiary positions used to formulate the half-angle rule with the discussion of the configuration space of such position changes in [18]. To my best knowledge, I have never seen such a derivation.

Without indicating the right or left eye in the binocular system, RVs ***r***^12^ corresponding to the eyes’ rotation when bifoveal fixation *F* ^1^ changes to *F* ^2^ in the stationary upright head,

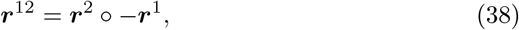

is the composition of RVs corresponding to the composition of rotation matrices *R*_2_(*ϕ*^2^, ***n***^2^)*R*_1_(−*ϕ*^1^, ***n***^1^). Here, RV ***r***^*i*^ = tan(*ϕ*^*i*^*/*2)***n***^*i*^ corresponds to the change *F*_*a*_ → *F*^*i*^ where *F*_*a*_ is the fixation of the ERP and *i* = 1, 2. For a full discussion, see [18].

Using (4) and (38), RVs ***r***^12^ is the following:

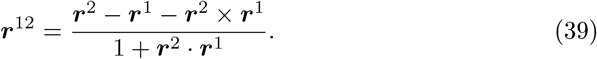

and from (1),

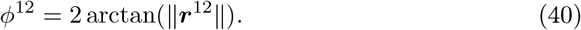

To obtain the angular velocity of the time-dependent RV ***r***(*t*), I introduce the notation for infinitesimally close fixation of *F*_1_ to *F*_2_. In the head frame ***r***^2^ − ***r***^1^ = *d****r***, is substituted into (39), denoting ***r***^12^ by *δ****r*** and dividing both sides by time *dt* elapsed between the fixations.

The rotation matrix (5) takes on the form *R* = *I* + *δ****r***× for infinitesimal RV *δ****r***, and (6) and (8) give *R****x*** − ***x*** = 2*δ****r*** × ***x***. Now, *d****x****/dt* = ***ω*** × ***x***, where ***ω*** is the instantaneous angular velocity vector in the head frame, i.e., ***ω***^*I*^. From the above discussion, we conclude that *δ****r*** = ^1^ ***ω***^*I*^*dt* and, hence,

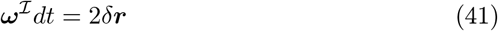

As *dt* approaches zero, substituting in (39) *δ****r*** for ***r***^12^, and *d****r*** for ***r***^2^ − ***r***^1^ and denoting *d****r****/dt* by 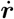, we have the angular velocity in the rotating eye’s frame,

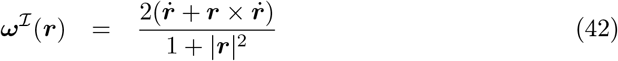

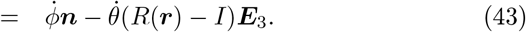

Of course, *δ****r***^12^*/dt* is not a derivative of any expression when *dt* → 0.

To obtain the angular velocity in the eye-rotating frame, I use Rodrigues’ identity

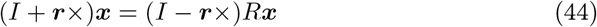

with a simple geometric presentation in Figure 2 in [35].

First, I note, using (44), that

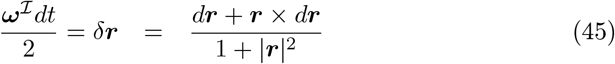

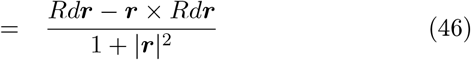

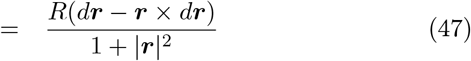

where I used that *R****r*** = ***r*** as the RV ***r*** is eigenvalue of *R*. We conclude that

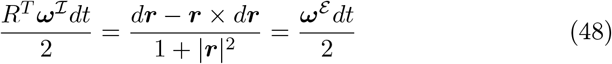

because *R*^*T*^ ***ω***^*I*^ = ***ω***^*E*^.

Thus, we obtain the angular velocity in the ERP initial frame

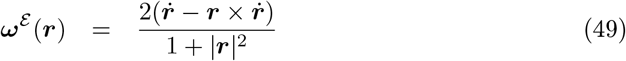

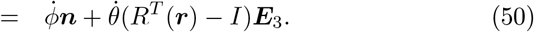

Similarly, we also easily obtain

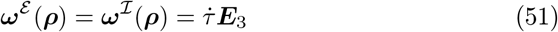

for ***ρ*** = tan(*τ/*2)***E***_3_.

The angular velocity for the composition of 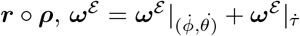 for the moving eye frame and 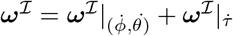 for the ERP initial frame can be obtained from (36) and (37) in the rotating eye frame and the ERP initial frame, respectively.

## 9 Discussion

Complementing the results in [18], which were motivated and demonstrated through GeoGebra simulations, I present a fully geometric formulation here. It includes a precise decomposition of the eye rotation into torsion-free and torsional components. By carefully introducing a parametrization of the eye’s geometry compatible with the ‘*z*-*x*-*z*’ original Euler-angle parametrization, the computed angular velocity correspondingly decomposes into torsion-free and torsional components. Further, following the results obtained in [18], a novel derivation of angular velocity is proposed. Future work will extend the presented geometric analysis to the binocular setting.

## Appendix

### A.1 Euler angle parametrization and ocular torsion

The ‘*z*-*x*-*z*’ parametrization of rotations introduced by Euler and frequently discussed in the literature, see, for example, [21], corresponds exactly to the rotation matrix (20) given by RVs ***r*** = tan(*ϕ/*2)***n*** and ***q*** = tan(*τ/*2)***E***_3_,

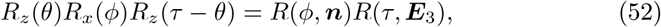

where all angles are time-dependent. We conclude that the vector ***n*** (denoted by ***n***_*r*_ and given explicitly in Fig. 2 (a) and obtained in (15) in terms of Euler’s angles) defines the line of nodes of this Euler parametrization of rotations. I recall that the line of nodes is defined as the intersection of the plane of the ERP spanned by ***E***_1_, ***E***_2_ and the rotated plane spanned by ***e***_1*r*_, ***e***_2*r*_ (cf. Fig. 2), which follows from the construction of ***n***_*r*_ for the right AE in (14).

The question is what ocular torsion corresponds to in the Euler parametrization for a rotated eye, where the angle *ψ* = *τ* − *θ* is referred to as *twist*, which for a spinning top represents rotation about its axis of symmetry. To understand the answer, I note that by identifying *R*(*τ*, ***E***_3_) with *R*_*z*_(*τ*) and applying to the right sides of expressions in (52), we obtain

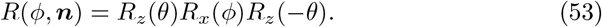

Thus, like *R*(*ϕ*, ***n***), *R*_*z*_(*θ*)*R*_*x*_(*ϕ*)*R*_*z*_(−*θ*) is a geodesic rotation, therefore, torsion-free. It shows that the real ocular torsion is indeed given by the angle *τ*.

### A.2 Angular velocity from RVs

Following [23], with the angular velocity in the eye rotating frame 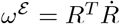, the angular velocity in the Euler-Rodrigues parameters *ρ* = cos(*φ/*2) and ***e*** = sin(*φ/*2)***n*** discussed in Sec. 3, the angular velocity becomes

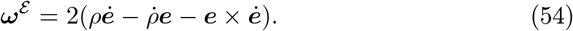

It can be further written in terms of Rodrigues’ vector ***s*** = ***e****/ρ*, as follows,

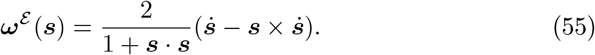

Similarly, the angular velocity in the ERP initial frame takes on the form,

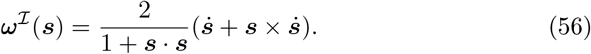

I compute first ***ω***^*E*^ (***s***) and then ***ω***^*I*^(***s***) when ***s*** is replaced with ***r*** = tan(*ϕ/*2)(cos *θ*, sin *θ*, 0)^*T*^ and next with ***q*** = tan(*τ/*2)(0, 0, 1)^*T*^. To do this, I first calculate 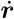 and 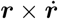. By differentiating ***r***,

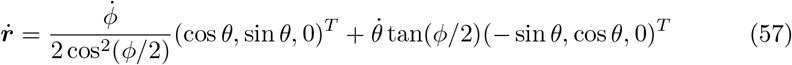

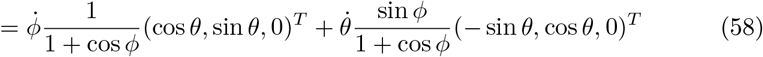

so that

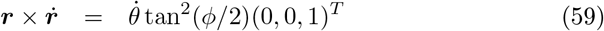

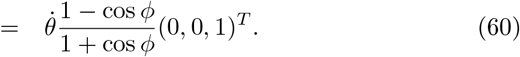

Substituting into,

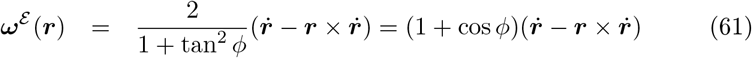

we obtain

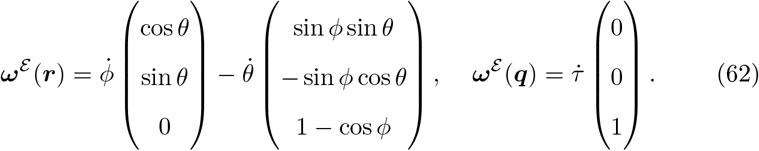

Similarly, we obtain

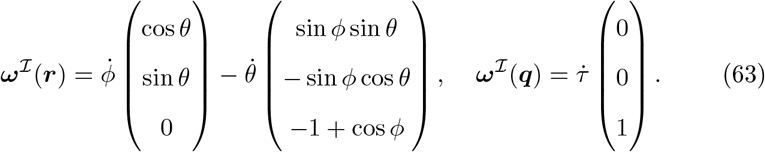

Using the relations (29), (30), and (31) from Sec. 7.2 we write

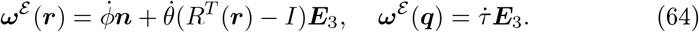

Similarly, we obtain

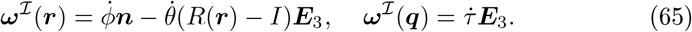

## Notes

### Competing Interest Statement

The authors have declared no competing interest.

### Summary of Updates

I improved the English throughout the paper. I rewrote Section 8 (New derivation of the angular velocity). It corrects the previous derivation.

